# Distinct Genetic Populations and Resistance Backgrounds of the Malaria Vector *Anopheles funestus* in Tanzania

**DOI:** 10.1101/2025.01.21.634154

**Authors:** Joel O. Odero, Ismail H. Nambunga, Hamis Bwanary, Gustav Mkandawile, John M. Paliga, Salum A. Mapua, Sophia H. Mwinyi, Halfan S. Ngowo, Nicodem J. Govella, Emmanuel W. Kaindoa, Frédéric Tripet, Anastasia Hernandez-Koutoucheva, Heather M. Ferguson, Chris S. Clarkson, Alistair Miles, David Weetman, Francesco Baldini, Fredros O. Okumu, Tristan P. W. Dennis

## Abstract

Population genetic analysis of mosquitoes is becoming increasingly important for understanding the distribution of insecticide resistance alleles, devising sustainable insecticide-based vector control approaches, and how malaria vector populations are structured in space. *Anopheles funestus* is the dominant malaria vector in Tanzania and most parts of East and Southern Africa. To better understand its population genomic structure in Tanzania, we sequenced the genomes of 334 individual *An. funestus* mosquitoes from 11 administrative regions. We found two genetically differentiated populations; one inland and at high altitude (found in Katavi, Kagera, Kigoma, and Mwanza) and a second coastal, at low altitude (found in Pwani, Morogoro, Tanga, Ruvuma, Mtwara, Dodoma, and Lindi), with differences in genetic diversity and inbreeding. We found asynchronous selective sweeps, associated with insecticide resistance phenotypes, at the *Cyp9k1* gene, and *Cyp6p* gene cluster, with distinct copy number-variant profiles between the coastal and inland populations. These results suggest that inland and coastal *An. funestus* populations have divergent histories, with the arid, central region of Tanzania, which also contains the Rift Valley being a possible barrier to gene flow. Such population disconnectedness should be considered for insecticide deployment, resistance management, and the rollout of novel genetic-based vector control approaches. These findings provide the most detailed study of Tanzanian *An. funestus* population structure and resistance genetics to date. Future research should examine the epidemiological relevance of this discontinuity in gene flow and whether these populations have different malaria transmission abilities.

## Introduction

*Anopheles funestus* sensu stricto, hereafter *Anopheles funestus,* is an increasingly important malaria vector in Eastern and Southern Africa, responsible for most malaria transmission in many localities (Msugupakulya et al., 2023). Despite its epidemiological significance, the species has been less studied than other major African vector species like *Anopheles gambiae*, primarily due to challenges in laboratory colonisation, unresolved taxonomic complexities and challenges in field observations of the species in its natural habitats (Kahamba et al., 2022; Ngowo, Hape, Matthiopoulos, Ferguson, & Okumu, 2021). The control of *An. funestus* is further complicated by widespread insecticide resistance (Coetzee & Koekemoer, 2013). Sustained use of insecticides in vector control, notably in insecticide-treated bed nets (ITN) and spraying indoor surfaces with insecticides (IRS), and in agricultural pest control has led to the evolution of insecticide-resistant phenotypes and reduced efficacy of these first-line vector control tools (Churcher, Lissenden, Griffin, Worrall, & Ranson, 2016; Hemingway et al., 2016). Consequently, the fight against malaria remains at a crossroads, with over 600,000 annual deaths, mostly in African children under 5 years old (W.H.O, 2024).

Genomic surveillance of malaria vectors is crucial for understanding critical aspects of their biology and forms the basis for developing targeted and effective control strategies. For example, population genomic analysis has revealed a new cryptic taxon within the *An. gambiae* complex, *gcx1* (Bissau molecular form), in West Africa (Caputo et al., 2024). Genomic analyses can also aid in predicting the spread of advantageous variants (Clarkson, Temple, & Miles, 2018) and the discovery of novel insecticide resistance mechanisms such as copy number variation (CNV), associated with pyrethroid and organophosphate resistance in *An. gambiae* (Lucas et al., 2019; Lucas et al., 2024). For instance, CNVs in the *Cyp6aa/Cyp6p* gene cluster (*Cyp6aap*_Dup33) are associated with deltamethrin resistance in *An. arabiensis* (Lucas et al., 2024). This is in addition to the recent discovery of knockdown resistance (*kdr*) genotypes associated with organochlorine (DDT) resistance phenotypes in *An. funestus* (Odero, Dennis, et al., 2024). Understanding the genetic structure and variation in mosquitoes is also crucial in estimating the viability of transgenic approaches for mosquito control such as gene drives (Wang et al., 2021). Recent advances in molecular techniques and the availability of reference genomes (Ayala et al., 2022) have spurred research towards understanding population diversity in the previously understudied malaria vector, *An. funestus*.

*Anopheles funestus* mediates 90% of malaria transmissions in parts of Tanzania (Mapua et al., 2022; Matowo et al., 2021) and is the leading vector even in locations where other malaria vectors (e.g. *An. arabiensis*) are more numerically dominant (Matowo et al., 2023; Mwalimu et al., 2024). The important role of *An. funestus* in malaria transmission is mostly replicated across eastern and southern African countries (Msugupakulya et al., 2023). *Anopheles funestus* has a wide ecological range spanning tropical and sub-tropical Africa (Odero et al., 2023). Previous studies of *An. funestus* shows broad-scale structuring into eastern, western, and central African genetic populations (Michel et al., 2005). Though studies investigating population diversity in *An. funestus* are limited, studies of *An. gambiae* indicates that geographical features (Lehmann et al., 2003), ecological variations and other genetic discontinuities play important roles in structuring *Anopheles* populations (Pinto et al., 2013). For instance, the Great Rift Valley has been hypothesised as a major gene-flow barrier in East and Southern Africa; as reflected by the presence of genetically distinct *An. funestus* populations on either side of the valley (Kamau, Hunt, & Coetzee, 2002; Michel et al., 2005). Continental mitochondrial analysis of *An. funestus* has discovered two distinct lineages on either side of the Rift Valley; a ubiquitous lineage I and an eastern and southern Africa-restricted lineage II (Jones et al., 2018). Due to restricted gene flow and adaptation, such lineages and diversity potentially impact the spread of beneficial genetic variation and constructs across populations.

Mutations in the major metabolic resistance genes, *Cyp6p9a* and *Cyp6p9b,* are widespread in pyrethroid-resistant *An. funestus* mosquitoes from East and Southern Africa but absent in West and Central Africa (Mugenzi et al., 2019; Weedall et al., 2019). Conversely, resistance to DDT is conferred by the glutathione S-transferase epsilon (*L119F-GSTe2*) mutation in *An. funestus* is largely restricted to West and Central Africa (Tchigossou et al., 2020) with evidence of spread to Eastern Africa (Odero, Nambunga, et al., 2024). These patterns depict restricted interaction and limited movement of these resistance alleles across the continental range of *An. funestus.* Our recent analysis of resistance markers within *An. funestus* populations in Tanzania, revealed an ongoing south-north directional spread of the *Cyp6p9a* and *Cyp6p9b* genotypes; suggesting landscape feature-driven gene-flow barriers or breakdown of past genetic barriers (Odero, Nambunga, et al., 2024). However, despite this spatial genetic variation, phenotypic resistance profiles were broadly similar in *An. funestus* from nationwide (Odero, Nambunga, et al., 2024). These findings raised the following questions pertinent to malaria control in Tanzania and beyond: i) how are Tanzanian *An. funestus* populations genetically structured, ii) what are the main drivers of genetic differences, and iii) how does genetic variation relate to patterns of phenotypic insecticide resistance and population size history?

To examine these questions, we analysed the whole genome sequences of 334 *An. funestus* mosquitoes sampled from 11 administrative regions with varying ecologies and malaria burden across mainland Tanzania. We found not only marked population structure within and between populations of *An. funestus,* potentially determined by the Rift Valley, also found that the distribution of metabolic resistance determinants varied according to population, suggesting divergent genomic architectures underlying a consistent resistance phenotype across Tanzania.

## Materials and Methods

### Population sampling and sequencing

*Anopheles funestus* specimens analysed in this study were collected from 11 locations spanning the eco-geography of Tanzania (**Figure 1A**). The collections formed part of a countrywide *An. funestus* surveillance project in Tanzania and were subsequently incorporated into the MalariaGEN *Anopheles funestus* genomic surveillance project database (https://www.malariagen.net/projects/anopheles-funestus-genomic-surveillance-project). Most mosquitoes were collected in households between 2021 and 2023 using CDC light traps and mechanical aspirators. They were sorted by sex and taxa and *An. funestus* group mosquitoes were preserved individually in 96-well plates containing 80% ethanol.

**Figure 1:**
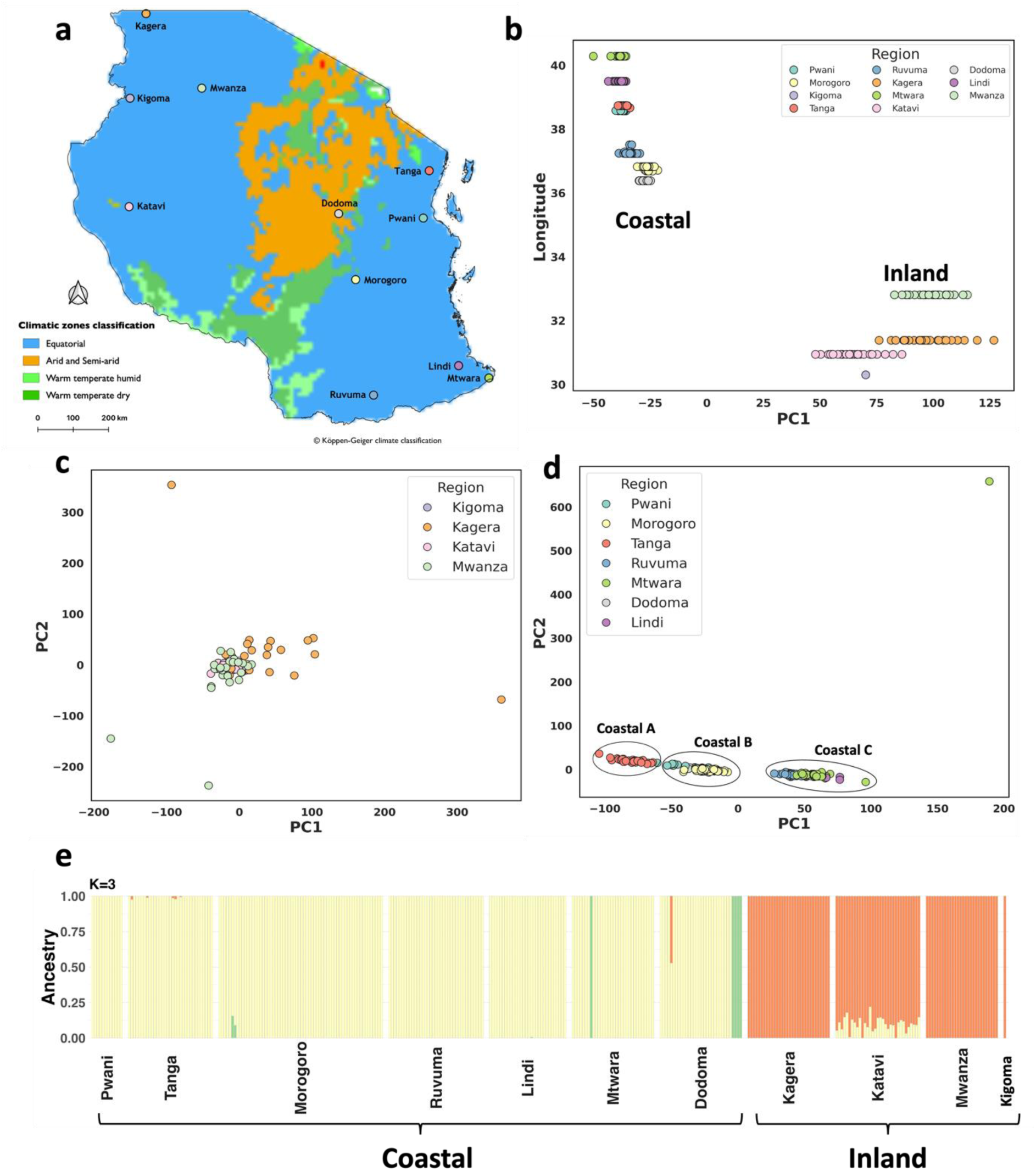
The population structure of *An. funestus* in Tanzania. (**A**) Map of *An. funestus* collection locations. Points indicate sample collection locations. The point colour indicates the administrative region from which samples were collected. (**B**) PCA plot of the first principal components and longitude from the 2RL chromosome. The colours denote the sampling location (regions). (**C**) PCA plot of the first two principal components from the 2RL chromosome for inland samples from the north and west; Kigoma, Kagera, and Mwanza. (**D**) PCA plot of the first two principal components from the 2RL chromosome for coastal samples from the south and east; Pwani, Morogoro, Tanga, Ruvuma, Mtwara, Dodoma, Katavi, and Lindi. (**E**) ADMIXTURE proportions: each mosquito is represented as a vertical bar on the y-axis, and the bar’s colour represents the proportion of the genome inherited from each of *K* = 3 inferred ancestral populations. The x-axis represents the 11 regions where the samples originated.

Genomic DNA was extracted from individual mosquitoes using DNeasy Blood and Tissue Kits (Qiagen, Germany). A single band confirmed the purity and integrity of 1% agarose gel and a minimum DNA concentration of 20 ng/μl on a Qubit® 2.0 fluorometer. Samples that passed quality control were individually commercially whole-genome-sequenced at 30X.

### Sequence analysis and SNP calling

The sequences were processed following pipelines developed by the Sanger Institute. Detailed specifications of the sequence analysis, SNP calling, and haplotype phasing pipelines are available from the MalariaGEN GitHub repository (https://github.com/malariagen/pipelines/). Briefly, reads were mapped to the *An. funestus* reference genome idAnoFuneDA-416_04 (Ayala et al., 2022) with Burrows-Wheeler Aligner (BWA) version v0.7.15. Indel realignment was performed using Genome Analysis Toolkit (GATK) version 3.7-0 RealignerTargetCreator and IndelRealigner. Single nucleotide polymorphisms were called using GATK version 3.7-0 UnifiedGenotyper. Genotypes were called for each sample independently, in genotyping mode, given all possible alleles at all genomic sites where the reference base was not “N”. The aligned sequences in BAM format were stored in the European Nucleotide Archive (ENA) under study number PRJEB2141.

High-quality SNPs and haplotypes were identified using BWA version 0.7.15 and GATK version 3.7-0. Quality control involved the removal of samples with low mean coverage, removing cross-contaminated samples, running PCA to identify and remove population outliers, and sex confirmation by calling the sex of all samples based on the modal coverage ratio between the X chromosome and the autosomal chromosome arm 3R. Full quality control methods are available on the MalariaGEN vector data user guide (https://malariagen.github.io/vector-data/ag3/methods.html). We used decision-tree filters that identify genomic sites where SNP calling and genotyping are likely to be less reliable. Genotypes at biallelic SNPs that passed the decision-tree site filtering process were phased into haplotypes using a combination of read-backed and statistical phasing. Read-backed phasing was performed for each sample using WhatsHap version 1.0 (https://whatshap.readthedocs.io/). Statistical phasing was then performed using SHAPEIT4 version 4.2.1 (https://odelaneau.github.io/shapeit4/).

### Population structure

To estimate geographic genetic population structuring in *An. funestus* across Tanzania, we used principal component analysis (PCA) and a model-based ADMIXTURE ancestry analysis. PCA analysis was conducted on Google Colab following procedures outlined in the malariagen_data (v14.0.0) python API (https://malariagen.github.io/malariagen-data-python/latest/Af1.html). The PCA was run on an inversion-free region of chromosome 2RL: 57,604,655 - 90,000,000 obtained from the full data set via random down-sampling to 100,000 SNPs and plotted using the seaborn python package (Waskom, 2021). ADMIXTURE analysis was performed to classify the individual mosquitoes of unknown ancestry into discrete populations. The analysis was performed using the software, ADMIXTURE v1.3.0 (Alexander, Novembre, & Lange, 2009) on 100,000 SNPs randomly down-sampled from chromosome 2RL with *K* (a priori clusters) values ranging from 1-10. The best value of *K* with the lowest cross-validation error (CVE) of 0.10027 was *K* = 3 followed closely by *K* = 4 (CVE 0.10352) and *K* = 2 (CVE 0.10373) (**Figure S2**). ADMIXTURE was run using identical parameters across 5 different random seeds, and results were examined visually between runs to ensure consistency. The R package Starmie (https://github.com/sa-lee/starmie) was used to summarise the results and the output visualised using ggplot2 in R (Wickham, 2009).

### Genetic diversity

Genetic diversity was estimated across the populations by computing Nucleotide diversity (π) and Tajima’s D statistics. The analyses were conducted on Google Colab following procedures outlined in the MalariaGEN Python package https://malariagen.github.io/malariagen-data-python/latest/Af1.html. We determined the signatures of recent inbreeding among individuals in the populations by estimating genome-wide runs of homozygosity (ROH). The ROH were inferred using a hidden Markov model implemented in *scikit-allel* (Miles et al., 2024). For more information on the model, see the supplementary material of The *Anopheles gambiae* 1000 Genomes Consortium, 2017 (Anopheles gambiae Genomes et al., 2017). Due to the difficulty of inferring short ROH, ROH < 100,000bp in length was discarded as per this publication.

### Measures of population differentiation

Genome-wide *Fst* measures were used to determine the relative extent to which allele frequencies differ between groups of individuals and the degree of differentiation between the sampled populations. *Fst* values were estimated across each chromosome using the *fst_gwss* function in the malariagen_data python API (https://malariagen.github.io/malariagen-data-python/latest/Af1.html) with a genomic window size of 5000 and a minimum cohort size of 10. We also searched for genome-wide signatures of selective sweeps using Garud’s H12 selection statistic (Garud, Messer, Buzbas, & Petrov, 2015). H12 selection scans were performed on *An. funestus* genotypes by collection region where sample *n>*10 and a genomic window size of 5000 using the *h12_gwss* function in the malariagen_data python API.

### Haplotype clustering

To investigate the haplotype structure in *An. funestus* around known insecticide resistance genes which can give evidence for recent selection, we performed haplotype hierarchical clustering on all 334 mosquito genomes at transcript LOC125764713_t1, 2RL:8,685,464 - 8,690,407, and LOC125764232_t1, X:8339269 - 8341975, using a Hamming distance matrix, inferred from phased *An. funestus* haplotypes, using *plot_haplotype_clustering* function in the malariagen_data python API.

### Amino acid variations

To identify potential nucleotide polymorphisms *Cyp6p* and *Cyp9k1* genes, we extracted single nucleotide polymorphism (SNPs) altering the amino acid of transcripts LOC125764713_t1 and LOC125764232_t1. We computed the allele frequency on the mosquito cohorts defined by the region and collection year. We filtered out variant alleles with a frequency lower than 5% (Anopheles gambiae Genomes et al., 2017). The resulting frequencies of non-synonymous SNPs were visualised in R software (R.Core.Team, 2020).

### CNV frequencies

Copy number variation (CNV) frequency analysis was conducted on CNV alleles previously identified in MalariaGEN Af1.1-1.3 data release. Full details of CNV calling and phasing are available on the MalariaGEN Python package (https://malariagen.github.io/malariagen-data-python/latest/Af1.html). The CNV frequencies were computed on the *Cyp6p* gene cluster (*Rp1*) (2RL: 8.6 - 8.8 Mbp) and *Cyp9k1* genes (X: 8.35 - 8.55 Mbp) using the *gene_cnv_frequencies* function in the malariagen_data python API.

## Results

### Population Structure

We estimated population structure using principal component analysis (PCA) and a model-based ADMIXTURE ancestry analysis (Alexander et al., 2009). To avoid the likely confounding effects caused by chromosomal inversion polymorphisms and recombination on the *An. funestus* genome (Boddé et al., 2024), our PCA analysis relied on an inversion-free section of chromosome 2RL (5.76 – 9 Mbp) and chromosome X (Boddé et al., 2024; Small et al., 2023). The PCA of chromosome 2RL revealed two main clusters (**Figure 1B & S1A**). The first cluster contained inland *An. funestus* populations from inland higher altitude regions of Katavi, Kigoma, Kagera, and Mwanza (hereafter referred to as the ‘Inland’ population). The second cluster contained coastal populations from lower altitude areas of Pwani, Morogoro, Tanga, Ruvuma, Mtwara, Dodoma, and Lindi (**Figure 1B**). The PCA based on chromosome X similarly showed the inland and coastal genetic clusters, but the samples from inland Tanzania were dispersed along PC1 and PC2 (**Figure S-1B**). We constructed separate PCA plots for samples from inland and coastal Tanzania to further investigate population structure within each major PCA grouping (as identified in **Figure 1B)**. The samples from inland Tanzania remained in a single cluster but with some dispersed samples from the Kagera region (**Figure 1C**). However, the second cluster containing coastal samples was separated into three genetic clusters with a clear pattern of geographic separation (**Figure 1D**). The first cluster contained samples from Tanga (coastal A), cluster two had Pwani, Dodoma, and Morogoro (coastal B), and cluster three Ruvuma, Mtwara, and Lindi (coastal C) (**Figure 1D**).

To understand the ancestral origin of the mosquitoes, ADMIXTURE analysis was performed to classify the individual mosquitoes into discrete populations using chromosome 2RL. The best ADMIXTURE value of *K* with the lowest cross-validation error (CVE) of 0.10027 was *K* = 3 **(Figure S-2).** The *K* = 3 disclosed two major groups of samples, with samples from inland Tanzania derived from two putative ancestral populations, and samples from coastal Tanzania derived from a third (**Figure 1E**).

### Genome-wide Population Diversity

To determine genome-wide population diversity, we computed nucleotide diversity (π) and Tajima’s D (Tajima, 1989) summary statistics using the SNP dataset from chromosome 2RL. This was calculated for populations with a minimum of 10 samples, hence samples from the Kigoma region in inland Tanzania were left out. The average nucleotide diversity (π) was higher in the inland samples (π = 0.01208) compared to the coastal populations (π = 0.01084) (**Figure 2A**). Similarly, the average values for Tajima’s D were lower in the inland samples (-1.21527) compared to the coastal populations (-0.47317) (**Figure 2B**). This suggested a greater genetic diversity and a relative excess of rare variants in inland *An. funestus* populations, in the northwestern regions (Kagera, Katavi, and Mwanza).

**Figure 2:**
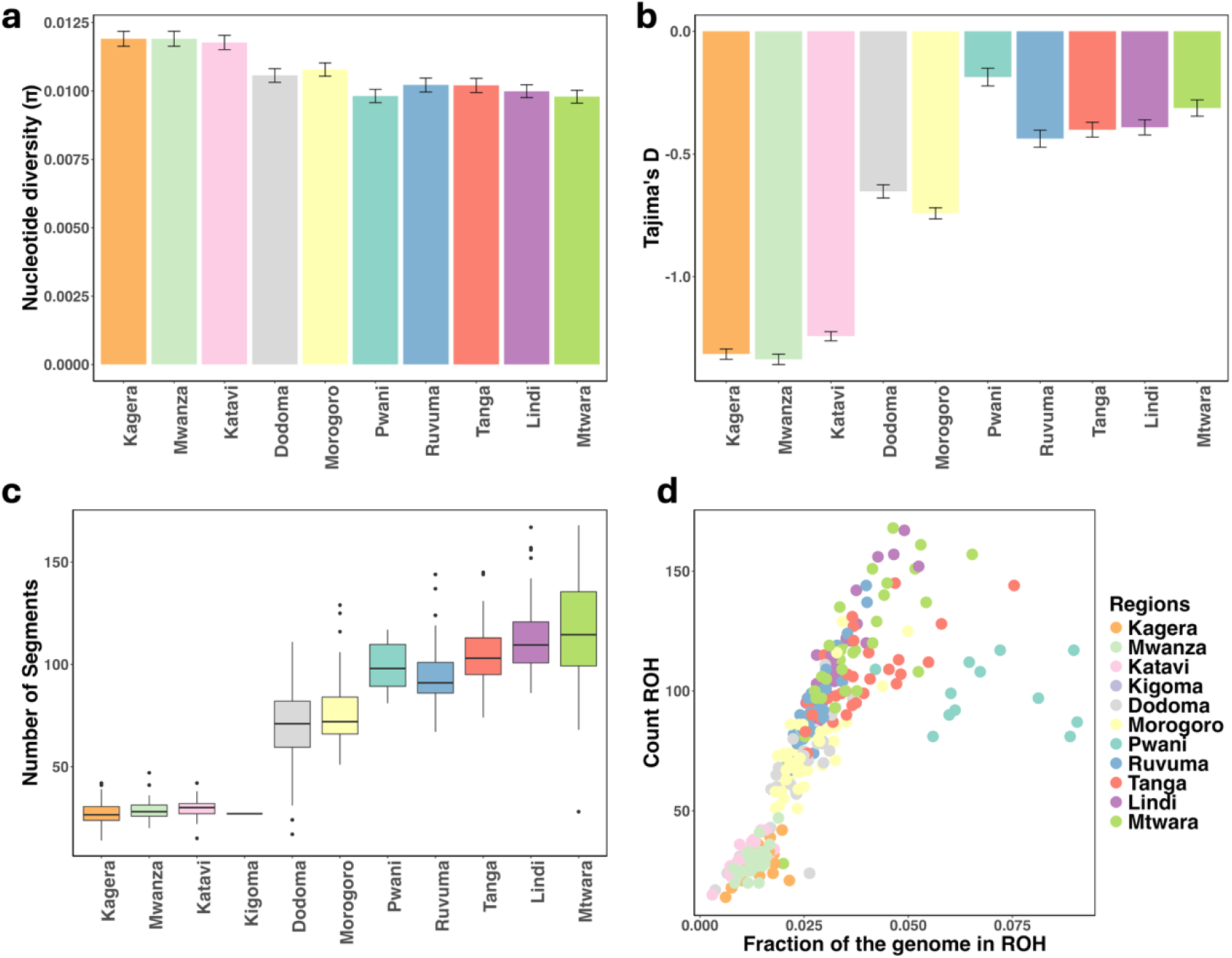
Genetic diversity in *An. funestus* populations in Tanzania. (**A**) Nucleotide diversity (*π*) was calculated using SNPs from chromosome 2RL. (**B**) Tajima’s D was calculated using SNPs from chromosome 2RL. (**C**) Boxplot showing the average number of segments of runs of homozygosity (ROH) per population across Tanzania. (**D**) Scatterplot showing the number of ROH (y-axis) and fraction of the genome in ROH (x-axis) in individual mosquitoes. Each marker represents a mosquito and is coloured by location (region).

The proportion of the individual genome in long runs of homozygosity (ROH) is informative of recent demographic processes (Ceballos, Joshi, Clark, Ramsay, & Wilson, 2018). Runs of homozygosity (ROH) are stretches of the genome where identical haplotypes are inherited from each parent due to inbreeding or recent common ancestry (Ceballos et al., 2018). Individuals from the inland populations (Kagera, Katavi, Kigoma, and Mwanza) generally had fewer ROH segments and a lower fraction of their genomes in ROH than from coastal (Dodoma, Morogoro, Pwani, Ruvuma, Tanga, Lindi, and Mtwara) which generally had more ROH segments and a higher fraction of their genomes in ROH, suggesting a history of recent inbreeding or a population bottleneck (**Figure 2C&D**).

### Genome-wide signatures of selection

We estimated between-population differentiation (*Fst*) and detected regions under possible recent selection (H12) across the whole mosquito genome (chromosome arms 2RL, 3RL, and X) (**Figure 3A**). Following the identification of genetic clustering (as shown in **Figure 1C&D)**, the analysis cohorts for pairwise *Fst* estimation and H12 were categorised into inland (found in samples from Katavi, Kigoma, Kagera, and Mwanza), coastal A (Tanga), coastal B (Morogoro, Pwani, and Dodoma), and coastal C (Ruvuma, Lindi, and Mtwara).

**Figure 3:**
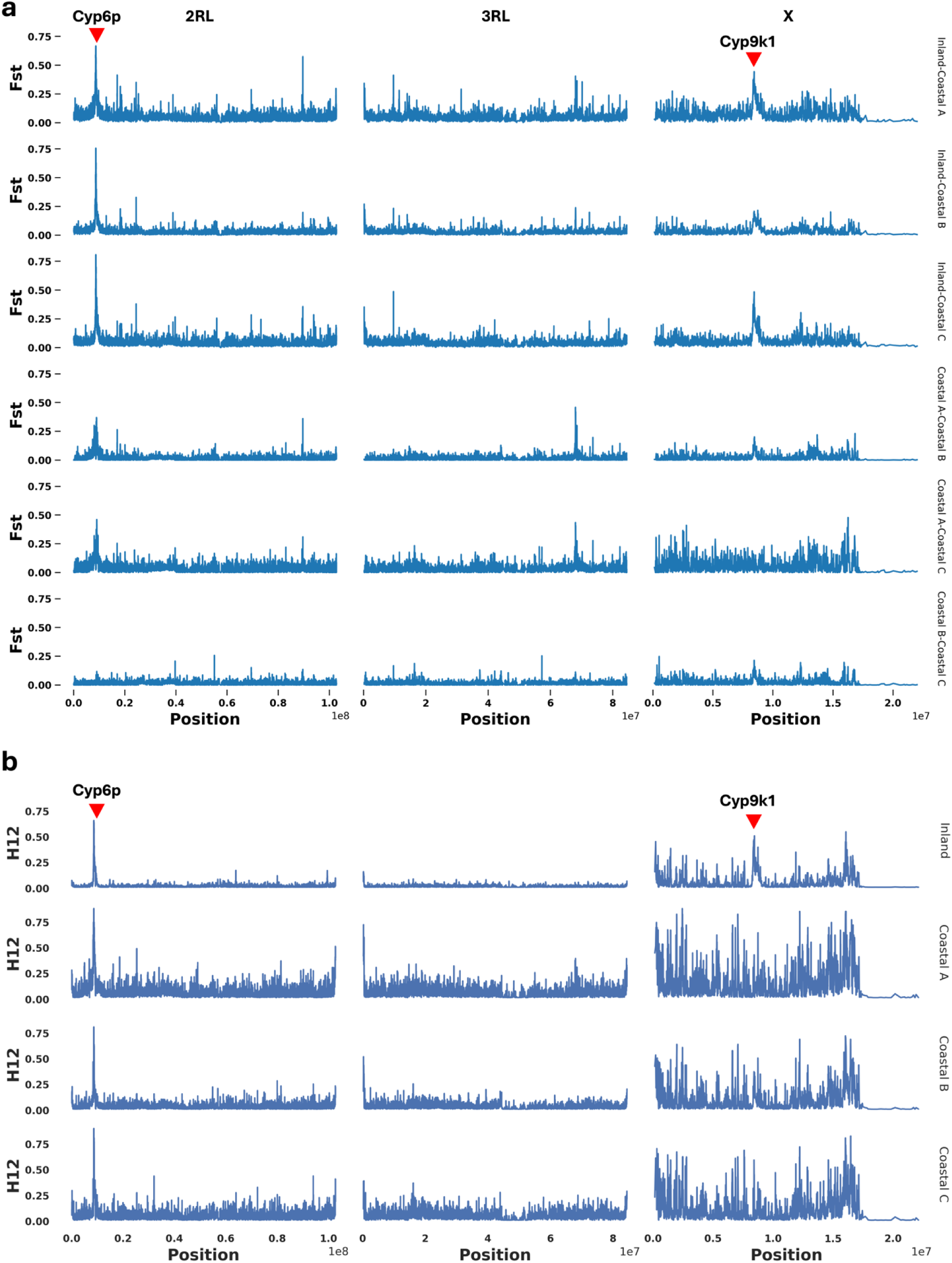
Genome-wide signatures of selection. (**A**) Genetic differentiation (*F_ST_*) at genomic windows along chromosomes 2RL, 3RL, and X. The X-axis indicates the position (in base pairs (bp)), and the Y-axis indicates the fixation index (*F_ST_*). (**B**) The measure of haplotype homozygosity (*H12*) genome-wide selection scans at genomic windows along chromosomes 2RL, 3RL, and X. The X-axis indicates the position (in base pairs (bp)), and the Y-axis indicates the selection statistic *H12*. The *H12* values range between 0 and 1, where zero indicates high haplotype diversity within a genomic window, and values closer to one indicate a strong selection of dominant haplotypes.

Genome-wide *Fst* measures the relative extent to which allele frequencies differ between groups of individuals and provides information about the degree of differentiation between the sampled populations. *Fst* values were generally higher between the inland and coastal A/B/C populations; with very high local peaks (*Fst* ∼0.8) between inland and coastal A/B/C around the genomic regions containing insecticide resistance genes, *Cyp6p* gene cluster on chromosome 2RL, and *Cyp9k1* on chromosome X, indicating strong differentiation at these regions (**Figure 3A**). Peaks in the *Cyp6p* cluster, and *Cyp9k1* were apparent between coastal A and B/C, though much smaller than the peaks between inland and coastal clusters in general (**Figure 3A**).

To detect regions of the genome that have undergone a recent selective sweep, we used a statistical measure of haplotype homozygosity (H12) (Garud et al., 2015). Genome-wide selection scans (GWSS) with the H12 statistic revealed a clear signal of elevated H12 at the *Cyp6* cluster across all populations (**Figure 3B**). *Cyp9k1* had a clear H12 elevation in the inland genetic cluster, but possible peaks of elevated H12 near *Cyp9k1* in the three coastal clusters were unclear against the relatively higher genome- wide H12 background (**Figure 3B**).

### Haplotype clustering around the Cyp6p9 and Cyp9k1 genes

To investigate the haplotype structure around known insecticide resistance genes with evidence for recent selection, we constructed haplotype clustering dendrograms from haplotypes in all 334 individual mosquitoes for the genes *Cyp6p9* (2RL:8,685,464 - 8,690,407) and *Cyp9k1* (X:8,448,477 - 8,450,887) (**Figure 4A&B**). The clustering dendrogram for the *Cyp6p* gene family revealed two major clades **(Figure 4A).** The first clade contained a swept haplotype in populations from the inland cluster (Katavi, Kigoma, Kagera, and Mwanza). The second clade was divided into three subclades containing haplotypes present in populations from coastal populations from the south and east of Tanzania **(Figure 4A).** The *Cyp9k1* dendrogram revealed three major clades **(Figure 4B).** The first clade contained a haplotype present only in samples from Mwanza in the inland populations. The second clade contained haplotypes that did not have geographic segregation based on sampling location **(Figure 4B).**

**Figure 4:**
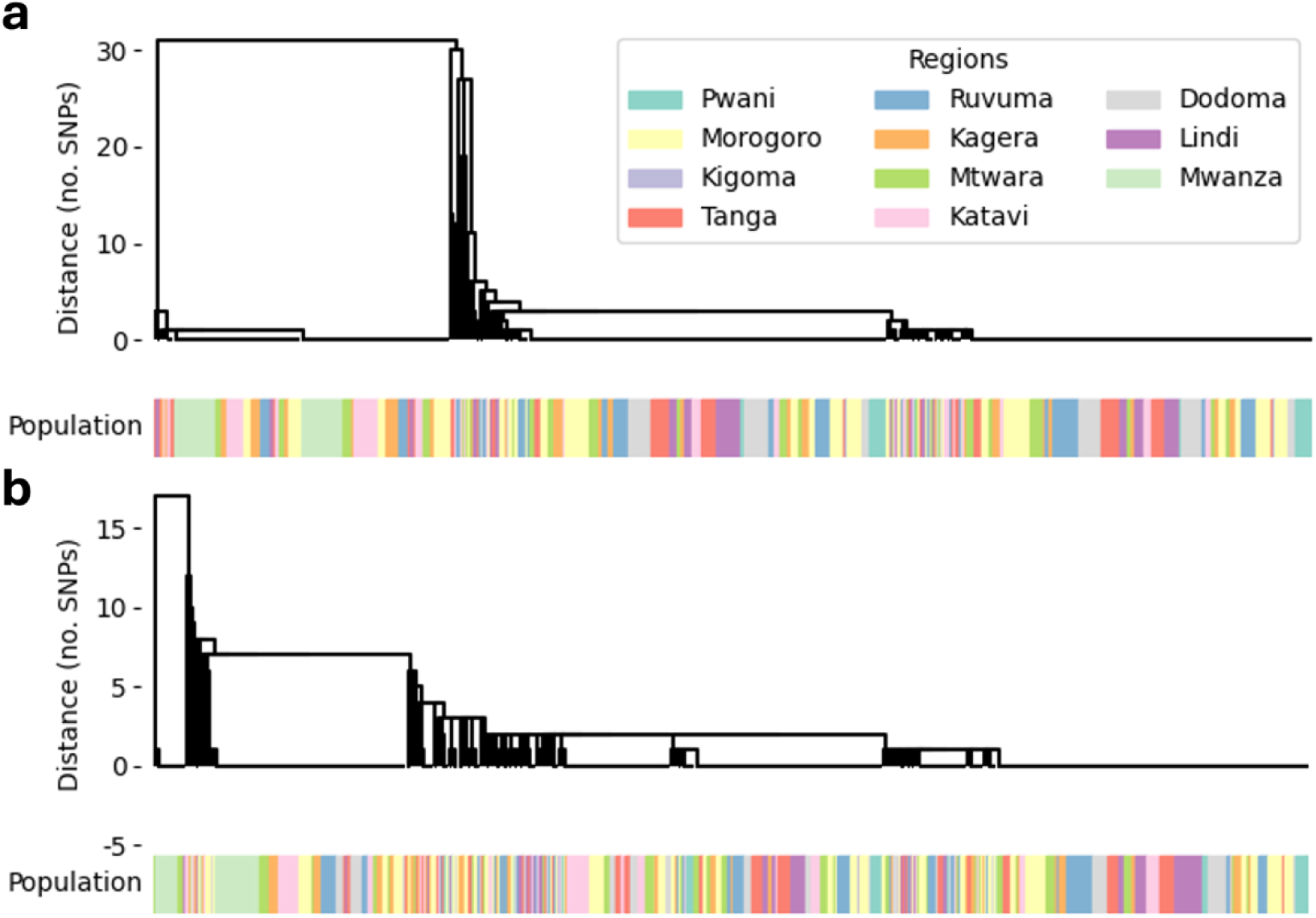
(A) Haplotype clustering over the *Cyp6p9* gene. (**B**) Haplotype clustering over *Cyp9k1* gene. The clusters are based on genetic distance as the number of SNPs and clusters with zero genetic distances indicate a selective sweep.

### Amino acid variation around the Cyp6p9 and Cyp9k1 genes

We investigated amino acid variation at the *Cyp6p* gene cluster (*Rp1*) (2RL: 8.6 - 8.8 Mbp) and *Cyp9k1* genes (X: 8.35 - 8.55 Mbp) by plotting non-synonymous amino acid variation with 5% minimum allele frequencies. We discovered 117 non-synonymous variants across the two (**Table S1**). Further analysis on *Rp1* was performed on transcript LOC125764713_t1 containing the *Cyp6p9a* which is crucial for insecticide detoxification in *An. funestus* populations from east and southern Africa (Mugenzi et al., 2019; Weedall et al., 2019). The *Cyp6p9a* gene had 13 non-synonymous amino acid variations with notably high frequencies of V103I (86 - 100%) and L122F (87 - 100%) across many populations (**Figure 5A**). However, both variants were absent in two inland populations (Kagera and Mwanza) and only at 32% frequency in Katavi (**Figure 5A**). Three additional variants, Y168H, V392F, and V359I, were present in the inland populations at greater than 40% but were absent in the coastal populations (**Figure 5A**). Further variants were detected in the *Cyp6p* gene in populations nationwide without a discernible geographical pattern (**Figure 5A**).

**Figure 5:**
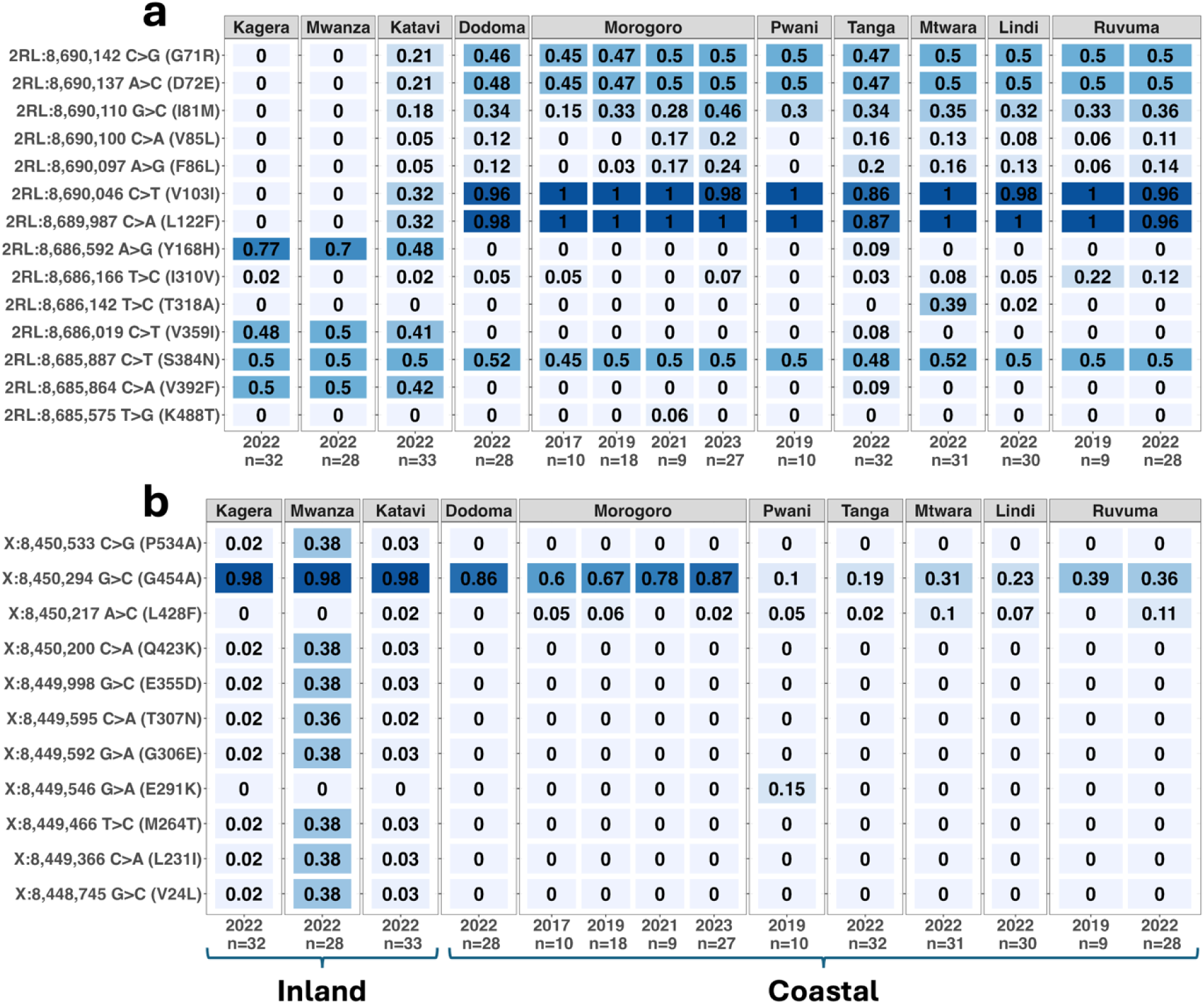
Amino acid variation around the *Cyp6p9* and *Cyp9k1* genes. (**A**) Heatmap of *Cyp6p9a* gene allele frequencies. (**B**) Heatmap of *Cyp9k1* gene allele frequencies. On both plots, the Y-axis labels indicate mutation effect, chromosome position, and nucleotide change. X-axis labels indicate the collection date and heatmap intensity indicates frequency where darker = higher, with frequency labelled in each heatmap facet.

The *Cyp9k1* gene, transcript LOC125764232_t1, had 11 amino acid variants, with variable frequencies of G454A across all populations (**Figure 5B**). The allele frequencies of G454A were at near fixation (98%) in samples collected from the inland cohort (Katavi, Kagera, and Mwanza) (**Figure 5B**). Two coastal B populations, Dodoma and Morogoro (2021), also had at least 80% frequency of G454A, with the coastal A (Tanga) and C (Ruvuma, Mtwara and Lindi) having between 10 - 36% (**Figure 5B**). In Morogoro, the frequency of the G454A mutation showed an increasing trend over 4 years, peaking at 87% in 2023, though this trend was insignificant (P = 0.06627, **Figure 5B**).

### Copy number variation around the Cyp6p gene cluster (*Rp1*) and Cyp9k1 genes

In addition to the target site (*kdr*) and metabolic resistance mechanisms, copy number variations (CNV) in the major resistance gene families have been demonstrated to increase resistance phenotypes in malaria vectors (Lucas et al., 2023). We investigated CNV frequencies of amplifications (increase in copy number) and deletions (reduction in copy number) at the *Cyp6p* gene cluster (*Rp1*) and *Cyp9k1* genes by a Hidden Markov Model (HMM). On the *Rp1*, CNVs around the probable cytochrome P450 6a13 gene (transcript LOC125764726) had the highest amplification frequency in Tanzanian populations (87 to 100%, **Figure 6**). The esterase E4-like gene amplification ranged from 30 - 70% with moderate CNV deletions in the inland populations (**Figure 6**). CNVs on the *Cyp6p9a* (transcript LOC125764713) were found in three coastal populations (Mtwara, Lindi and Ruvuma) (**Figure 6**). However, we did not find evidence of CNV amplification/deletion on the *Cyp9k1* gene within the populations (**Figure S3**).

**Figure 6:**
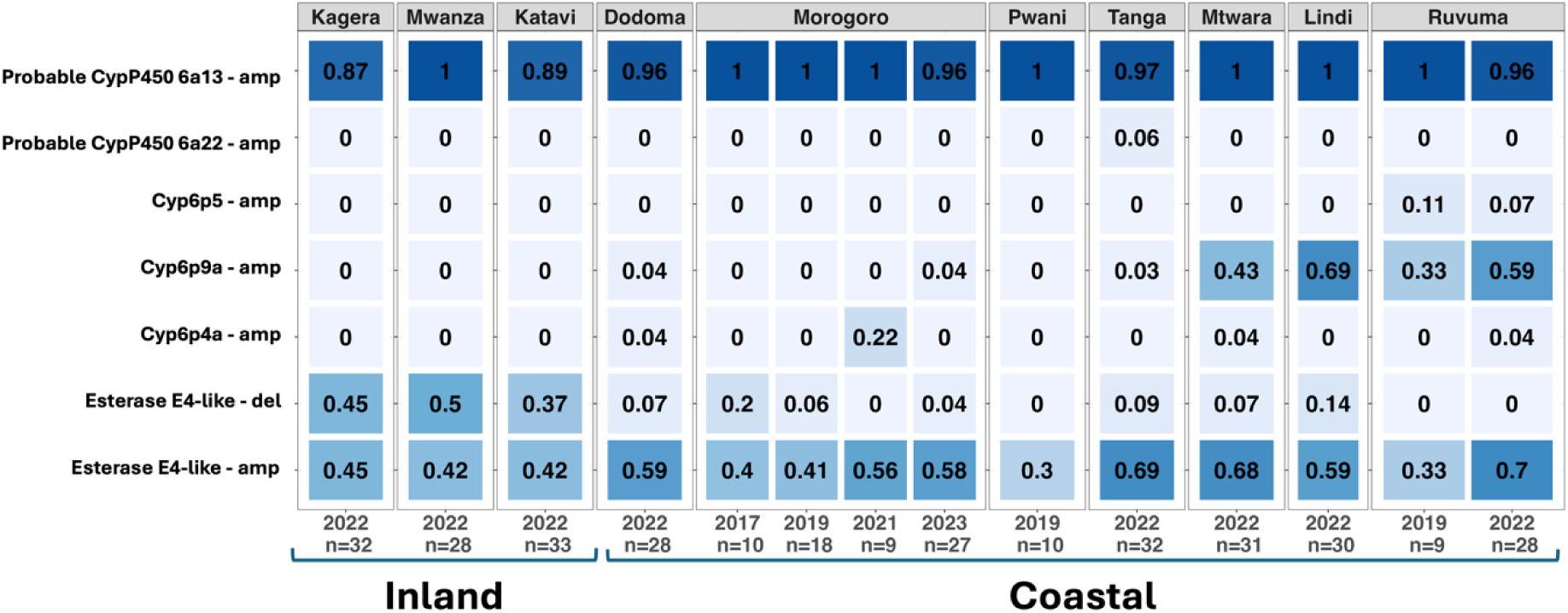
Copy number variations (CNV) in the major resistance gene family, *Cyp6p*, across Tanzania. CNV amplification is indicated by ‘amp’ and deletion by ‘del’. frequencies

## Discussion

Our findings demonstrate the existence of two genetically differentiated *An. funestus* populations in Tanzania; an inland population (comprising Katavi, Kagera, Kigoma, and Mwanza) and a geographically structured coastal population (found in the lower altitude coastal regions including Pwani, Morogoro, Tanga, Ruvuma, Mtwara, Dodoma, and Lindi). Genome-wide selection scans suggest that the population partitioning could be due to a combination of asynchronous selective sweeps at insecticide resistance genes, against a backdrop of possible demographic differences (e.g. population size or migration from neighbouring populations), evidenced by genome-wide differentiation, and distinct signatures of genetic diversity and recent inbreeding, between the two major populations. Recent population bottlenecks and inbreeding substantially reduce the frequency of rare genetic variants and overall genetic diversity. The inland *An. funestus* population had higher diversity, a lower Tajima’s D, and fewer ROH than the coastal population. This difference could be due to greater connectivity to continental *An. funestus* populations leading to a greater population size, or perhaps a greater effect of vector control on the comparatively geographically isolated coastal *An. funestus* population.

Inland and coastal populations of *An. funestus* in Tanzania are also located on either side of the eastern arm of the Rift Valley; making this landscape feature a plausible explanation for their observed differentiation. Major geographical features such as the Great Rift Valley have been postulated as a gene flow barrier across the east and southern African region (Lehmann et al., 1999; Michel et al., 2005). For example, *An. funestus* populations from the north of Malawi were genetically differentiated from those in the south; with the Rift Valley running between these sites hypothesised as the potential driver of this separation (Barnes et al., 2017). A similar association between the Rift Valley and the genetic structure of *An. funestus* populations in Kenya have been found (Ogola, Odero, Mwangangi, Masiga, & Tchouassi, 2019). Ecological barriers to gene flow such as the Congo Basin tropical rainforest have also been associated with population structuring in *An. coluzzii* populations across Africa (Pinto et al., 2013). Tanzania’s climate is largely equatorial with an arid desert cutting across the central part of the country (Beck-Johnson et al., 2013). These harsh arid conditions in central Tanzania are potentially less favourable to *An. funestus* and may thus also influence their genetic structuring. Indeed, vector surveillance data in Tanzania reveals that *An. arabiensis* is the predominant mosquito species in this arid desert region, while *An. funestus* and *An. gambiae* are more prevalent in the northwestern and southeastern zones of the country (Mwalimu et al., 2024). The potential impact of such geographic and climatic stratifications on local vector adaptation and genetic differentiation in Tanzania should be investigated further.

The strongest differentiation between the two populations was around genome regions containing metabolic resistance genes (*Cyp9k1* and the *Cyp6p* cluster); indicating these genes are undergoing asynchronous selective sweeps. For instance, analysis of the *Cyp6p9* gene demonstrated that V103I and L122F mutations were widespread and at high frequencies in the coastal *An. funestus* populations while the inland populations had Y168H, V392F, and V359I. However, these mutations have less impact on insecticide resistance phenotypes in this vector across Tanzania (Odero, Nambunga, et al., 2024). The selective sweeps in the *Cyp6p9* gene coincided with the patterns of distribution of key metabolic resistance alleles in Tanzania with *Cyp6p9a* and *Cyp6p9b* observed as fixed in the coastal populations of the south and east but either at low frequencies or undetectable in the inland populations of north and west Tanzania (Odero, Nambunga, et al., 2024). The structured allelic variations around genomic regions containing crucial metabolic resistance genes in *An. funestus* suggests underlying differences in the molecular basis of insecticide resistance across Tanzania. The cytochrome P450 G454A-*Cyp9k1* mutation was near fixation in the inland populations but present only at low frequencies in the coastal populations. This mutation has been demonstrated to confer phenotypic resistance to type I (permethrin) and II (deltamethrin) pyrethroids, commonly used on ITNs (Djoko Tagne et al., 2025; Hearn et al., 2022). Continentally, G454A-*Cyp9k1* is widespread in central Africa (Cameroon) and East Africa including Uganda (Djoko Tagne et al., 2025). We speculate that the presence of G454A-*Cyp9k1* at near fixation in inland Tanzania indicates that these populations are more connected to the larger African continent and coastal connected more to the southern African region where this mutation is either absent or at low frequencies (Djoko Tagne et al., 2025).

Our findings provide useful information when planning the implementation of transgenic vector control strategies such as gene drives and sterile insect releases in Tanzania. The divergence between the inland and coastal *An. funestus* populations with limited gene flow between them indicate that multiple strategic release sites across the potential genetic barriers would be required for nationwide implementation. However, in this study, the sampling locations were not uniformly spread across the country, leaving an area of ∼600 km between Dodoma in central Tanzania and Mwanza in the northwest unsampled. Isolation by distance (IBD) can be expected to give rise to genetic structuring between populations separated by such large geographical distances (van Strien, Holderegger, & Van Heck, 2015).

Follow-up studies should focus on sampling and sequencing from these areas with a collective outcome of a finer-scale genetic structure of this important malaria vector. Future analysis including *An. funestus* samples from countries neighbouring Tanzania (Boddé et al., 2024) will put the genetic variation and structure into a broader perspective. Such analysis will make it possible to discern the genetic connectedness between coastal Tanzania and continental Africa *An. funestus* populations and help pinpoint whether the observed structuring is best explained by the Rift Valley, or other ecological factors acting at landscape scales, and how this influences genetic diversity.

## Conclusion

Overall, this study provides a comprehensive population genomic description of *An. funestus* in Tanzania. We demonstrate that the vector is differentiated into two genetically distinct populations with historical and contemporary gene flow patterns. We also reveal directional sweeps of haplotypes, especially in the *Cyp6* gene cluster, and geographic allelic variations in the genomic regions around the insecticide resistance genes *Cyp6p* and *Cyp9k1*. The genomic dataset generated in this study, accessible through the European Nucleotide Archive under study number PRJEB2141, will be crucial to expanding capabilities for molecular surveillance of insecticide resistance in *An. funestus* Tanzania.

## Data availability

The scripts and Jupyter Notebook used to analyse genotype and haplotype data, and produce figures and tables are available from the GitHub repository: https://github.com/joelodero/TZ_funestus_genomics_2025. The sequencing data generated in this study have been deposited in the European Nucleotide Archive (https://www.ebi.ac.uk/ena/browser/home) under study number PRJEB2141.

## Acknowledgement

The authors express their sincere gratitude to the local authorities in Tanzania, community leaders, and residents of all study villages for their invaluable cooperation and permission to conduct this research. Ethical approvals for this project were obtained from the Ifakara Health Institute’s Institutional Review Board (Protocol ID: IHI/IRB/no: 26-2020) and the Medical Research Coordinating Committee (MRCC) at the National Institute for Medical Research (Protocol ID: NIMR/HQ/R.8a/Vol.IX/3495). This study was supported by the MalariaGEN Vector Observatory (https://www.malariagen.net/vobs/), an international collaboration working to build capacity for malaria vector genomic research and surveillance.

## Author contributions

The project was conceived and supervised by F.O.O., F.B., and T.P.W.D. Field mosquito collection was performed by J.O.O., I.H.N., H.B., J.P and G.M. Laboratory analysis, data acquisition and management, and preparing samples for whole-genome sequencing were performed by J.O.O. S.N.P. calling, and haplotype phasing were performed by A.H-K., A.M, and C.S.C. J.O.O. and T.P.W.D. led the data analysis and generated all figures and tables. The manuscript was drafted by J.O.O. and revised by all authors. Throughout the project, all authors have contributed key ideas that have shaped the work and the final paper.

## Supplementary

**Figure S1:**
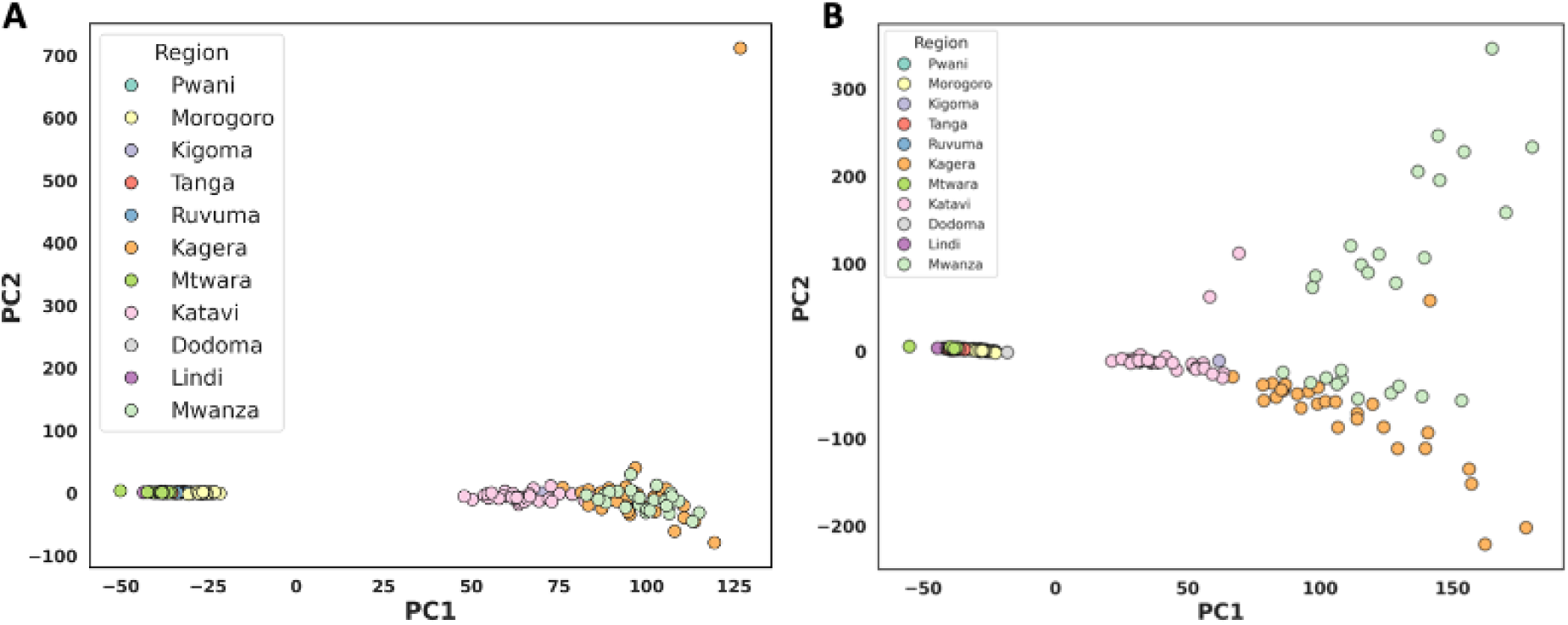
The population structure of *An. funestus* in Tanzania. (A) PCA plot of the first two principal components from the 2RL chromosome (B) PCA plot of the first two principal components from the X chromosome. The colours denote the sampling location (regions).

**Figure S2:**
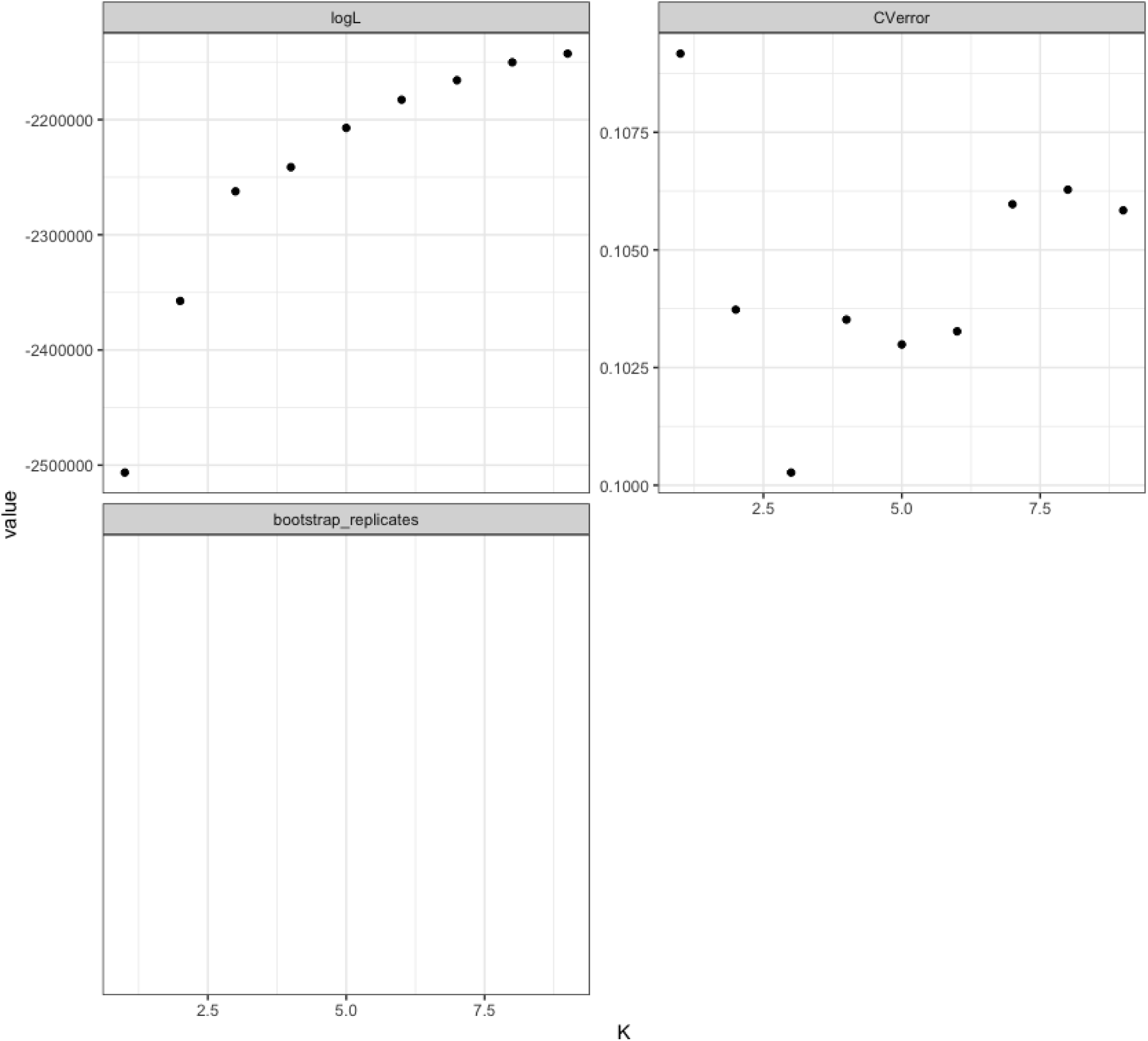
Validation of the best ADMIXTURE value of *K* with the lowest cross-validation error (CVE).

**Figure S3:**
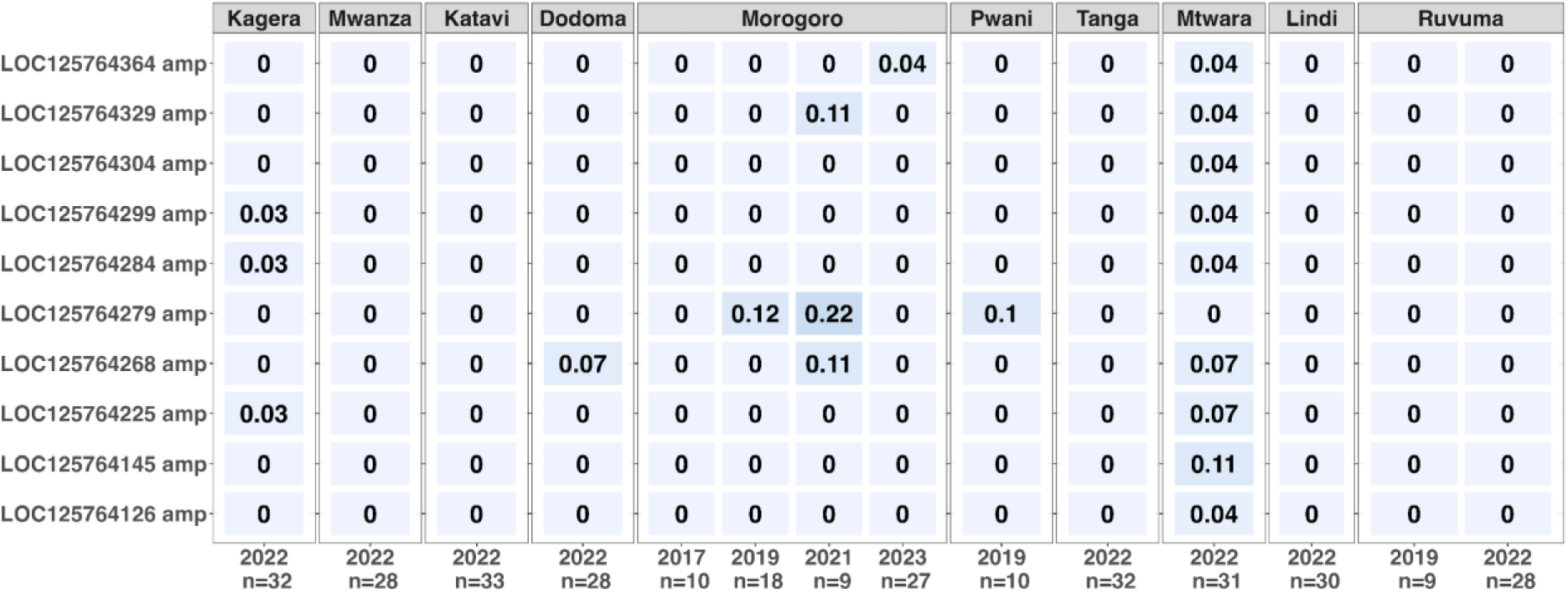
Copy number variations (CNV) in the major resistance gene family, *Cyp9k1*, across Tanzania. CNV amplification is indicated by ‘amp’ and deletion by ‘del’. frequencies

## Supplementary Table S1

https://docs.google.com/spreadsheets/d/1hr-qf35wCGZd-4JJfFOaE6hiQVuR4rSZ-ZBMVcIGk08/edit?gid=382865028#gid=382865028

